# A comment on priors for Bayesian occupancy models

**DOI:** 10.1101/138735

**Authors:** Joseph M. Northrup, Brian D. Gerber

## Abstract

Understanding patterns of species occurrence and the processes underlying these patterns is fundamental to the study of ecology. One of the more commonly used approaches to investigate species occurrence patterns is occupancy modeling, which can account for imperfect detection of a species during surveys. In recent years, there has been a proliferation of Bayesian modeling in ecology, which includes fitting Bayesian occupancy models. The Bayesian framework is appealing to ecologists for many reasons, including the ability to incorporate prior information through the specification of prior distributions on parameters. While ecologists often intend to choose priors so that they are “uninformative”; or “;vague”;, such priors can easily be unintentionally highly informative. Here we report on how the specification of a “vague” normally distributed (i.e., Gaussian) prior on coefficients in Bayesian occupancy models can unintentionally influence parameter estimation. Using both simulation and empirical examples, we illustrate how this issue likely compromises inference about species-habitat relationships. While the occurrence of this issue likely depends on the data set, researchers fitting Bayesian occupancy models should conduct sensitivity analyses to ensure intended inference, or use priors other than those most commonly used in the literature. We provide further suggestions for addressing this issue in occupancy studies, and an online tool for exploring this issue under different contexts.

## Introduction

Understanding species distributions, and the environmental factors that influence occurrence is fundamental to ecology. Our knowledge of many well-studied topics in ecology, including niche partitioning, trophic interactions and metapopulation dynamics, depend on knowing which species occur in an area and why. Furthermore, occurrence patterns are critical when making conservation and management decisions; placement of reserve boundaries, or assessments of whether development will impact threatened and endangered species depend entirely on knowing whether a target species is present. Research on the patterns and drivers of species occurrence has been ongoing for many years (see Guisan and Thuiller 2005 for a brief discussion), with major advancements, especially over the past two decades (see Elith and Leathwick 2009 for a review). These advancements have stemmed from a combination of enhanced computational power, the advent of geographical information systems (GIS), and the development of a diversity of field-sampling and statistical modeling approaches that allow for detailed assessments of species habitat-relationships and the ensuing distribution patterns.

At the forefront of the methodological advancements in modeling species distributions is the explicit recognition and correction for sampling biases, such as the non-detection of a species in an area, despite it being present (i.e., false-negatives; MacKenzie et al. 2002). These ‘occupancy models’ can account for the inherent imperfect detection of a species by simultaneously modeling the observation and occurrence processes. The development and refinement of these types of models has been a major focus of the ecological literature; there are numerous publications developing and describing occupancy models designed to address different ecological processes or sampling designs (see Kéry and Royle 2015), along with books acting as “how-to” guides (Royle and Dorazio 2008, Kéry and Royle 2015) and software for readily fitting occupancy models to ecological data (e.g. unmarked; Fiske and Chandler 2011; MARK; White and Burnham 1999; PRESENCE; Hines 2006). These resources have allowed researchers to apply occupancy models to a range of ecological questions.

Concomitant with the increasing prevalence of occupancy models has been the increase in the use of Bayesian statistics in ecology (Clark 2005, McCarthy 2007, Hobbs and Hooten 2015). The adoption of Bayesian statistics by ecologists has likely been driven by a number of factors, including the straightforward manner in which hierarchical or multi-level models can be specified and fit. Occupancy models are naturally structured hierarchically (see model below) making them straightforward to fit using Bayesian methods and there are numerous published and online resources that provide code to do so (e.g. Royle and Dorazio 2008, Kéry and Royle 2015). The increase in the availability of these resources has made the application of Bayesian methods more approachable for practitioners and researchers.

A potential risk of the proliferation of easily accessible software and code is that researchers are perhaps fitting models without a clear understanding of the consequences of modeling choices. In Bayesian modeling, one choice that has the potential to strongly influence statistical inference is that of prior distributions (Gelman et al. 2014). Briefly, Bayesian inference focuses on summarizing posterior distributions of model parameters, which are informed jointly by the likelihood and the prior distributions. The relative influence each has on the posterior distribution depends on their information quantity. A model’s likelihood is determined entirely by the data, while prior distributions represent our best knowledge about the distribution of a parameter prior to model fitting. Guidance on the choice of priors when fitting Bayesian models in ecology is limited. In our experience, researchers typically attempt to choose priors such that they are expected to have minimal or no influence on the resulting inference (i.e., “flat,” “vague,” or “uninformative” priors). Researchers commonly pick these priors so that parametric inference is primarily driven by the data, rather than the prior (e.g. Jones et al. 2011). However, seemingly uninformative priors often can have strong unintended consequences (Lele and Dennis 2009, Seaman III et al. 2012). Here we explore this issue with a specific focus on occupancy models. We show how under certain conditions a commonly used prior can strongly influence statistical inference. We provide examples of when the use of this prior is an issue and offer guidance when fitting Bayesian occupancy models.

### A basic Bayesian occupancy model

In the analyses and discussion below we focus on a simple site occupancy model, formulated in a hierarchical Bayesian framework, which takes the following form,

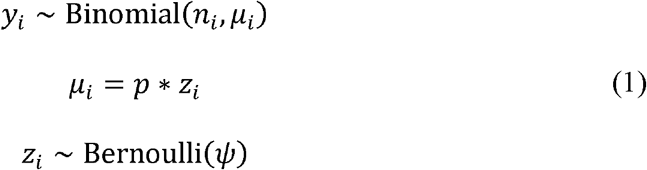

where *y*_*i*_ indicates the number of detections at site *i*, out of a total of *n*_*i*_ sampling occasions, *z*_*i*_ is a latent (unobserved) parameter indicating the true occupancy state of the site (1 = occupied and 0 = unoccupied), *p* is the probability of detecting a species at the site conditional on it being occupied, and ψ is the probability that a site is occupied. In a full Bayesian analysis, prior distributions would be specified for the unknown parameters, *p* and *ψ*. Often, uniform prior distributions between 0 and 1 are chosen (e.g. Kéry and Royle 2015).

Typically, researchers are interested in investigating hypotheses of whether environmental covariates influence species occurrence. In this case, the above model can be extended to include covariates on the occupancy process, which is most often specified as a logit regression model as

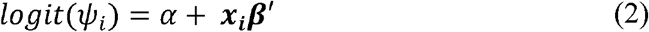

where *ψ*_*i*_ is the site-specific probability of occupancy, which is influenced by a matrix of covariates ***x***_*i*_ (for example land cover type at the site), corresponding vector of coefficients ***β*** and an intercept, *α*. The logit is a link function (i.e., 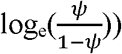) that takes probability values, which are restricted between 0 and 1, and projects them to values on the real number line, making estimation easier due to the lack of numerical boundaries. To recover occupancy probabilities, we use the inverse-logit of the linear combination of the intercept and covariates (i.e., logit 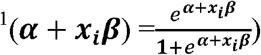. We note that other link functions are available for this model, such as the probit or complementary log-log link, but in our experience, these link functions are rarely used in the ecological literature.

### The issue of normally distributed priors in occupancy modeling

The convention in Bayesian regression models is to specify normally distributed (i.e., Gaussian distribution) priors for the intercept (*α*) and coefficients (***β***), with a mean of 0 and some standard deviation ( σ; e.g., α ~ Normal(0, σ); Gelman and Hill 2007; technically, in the example above a multivariate Normal prior with a vector of 0s for the mean and a covariance matrix with 0’s in all the off-diagonal positions would be used for ***β***). Normally distributed priors are a sensible choice for a range of reasons discussed elsewhere (e.g. Gelman and Hill 2007), but see Gelman et al. (2008) for a discussion of why alternative priors might be preferred in certain cases. When conducting regression analyses, researchers typically specify these priors with a large standard deviation (*σ*) hoping to diffuse the influence of the prior. In simple linear regression, this choice of prior has the intended result. However, the use of such “vague” Normal priors in occupancy modeling has a potentially pernicious outcome. The issue is that the logit transformation is non-linear such that as values become more negative or more positive, the transformed probability values approach zero and one, respectively (Fig. S1). This nonlinearity in the transformation leads to some priors that are intended to be “uninformative” becoming informative on the probability scale and strongly bimodal with large values of σ (Fig. 1 and Fig. S2). It is a mathematical truism that a normally distributed prior is not invariant to this transformation—however, we believe that the consequences for modeling occupancy (as well as any logistic regression model) are not well appreciated in the ecological literature. Below, we demonstrate the influence of this assumed prior distribution on inference from occupancy models, using both simulated and empirical datasets.

**Figure 1.**
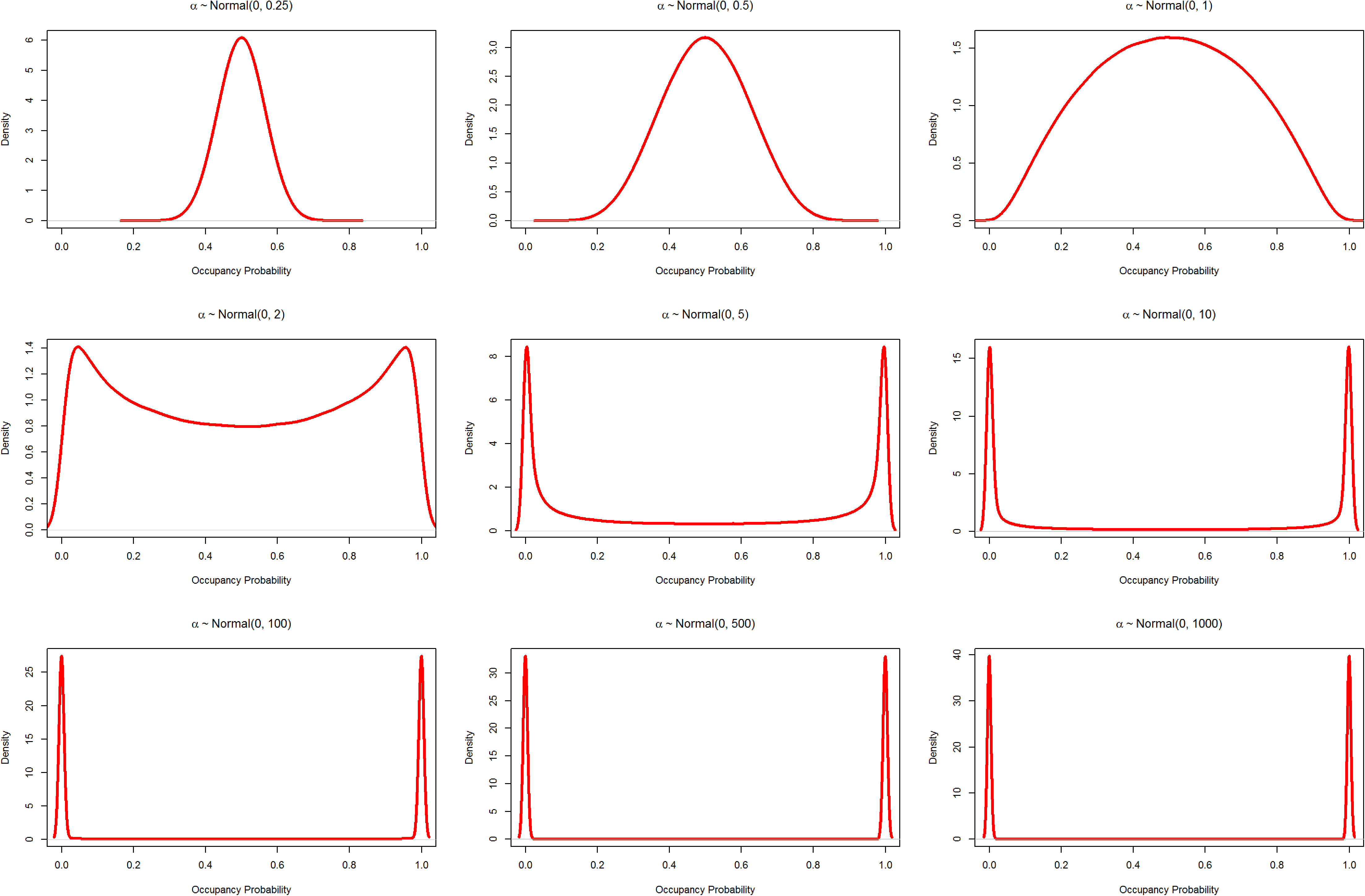
Demonstration of a Normal prior distribution transformed to the probability scale (*α* is the occupancy probability before transformation to the probability scale using the logit link; see equation 2 in the text); panels vary by the standard deviation (*σ*) of the prior distribution. A small *σ* gives high probability density around zero, while increasing levels move this probability density towards zero and one, which eventually begins to accumulate near these values. Note that y-axes differ substantially among the panels.

## Methods

We first demonstrate the influence of a logit-Normal prior by simulating example occupancy datasets (using *ψ* = 0.9, *p* = 0.2, *n* = 10, where *n* is the number of occasions) at a varying number of sites (50, 100, 200, and 400). For each dataset, we first fit the data in a maximum likelihood framework using the statistical program MARK (White and Burnham 1999) via the R package ‘RMark’ (Laake 2013) in the programming language R (R Core Team 2015). Next, to illustrate the influence of the prior relative to the likelihood, we fit these models in a Bayesian framework (above model without covariates; *logit*(*ψ*_*i*_) = *α* and prior *α* ~ *Normal*(0,τ)) using JAGS (Plummer 2015a) via the ‘rjags’ package (Plummer 2015b; see Appendix S1 and S2 for example JAGS code). In JAGS, the uncertainty parameter for the Normal distribution is specified as the precision(τ), which is 1/*σ*^2^, where *σ* is the standard deviation of the Normaldistribution. We fit the Bayesian model with normally distributed priors with a values of 0.25, 0.5, 1, 2, 5, 10, 100, 500, and 1000; algorithms were run for 10,000 Markov chain Monte Carlo (MCMC) iterations, removing the first 5,000 as a period of burn-in. We investigated convergence in both paradigms by fitting the models with random initial values, checking for estimate consistency. The parameters from the Bayesian analysis were also investigated for convergence by visually examining posterior distribution trace plots to ensure proper mixing and by calculating the Gelman-Rubin diagnostic (Gelman and Rubin 1992) to ensure values were close to 1, which they always were. We compare the likelihood results, which are not influenced by the assumed prior distribution, with the Bayesian results by plotting posterior distributions of *ψ* for each dataset and prior, along with the maximum likelihood estimate (MLE). Assuming convergence and a sufficiently large number of samples, the discrepancy between the posterior mode (i.e., most probable value) and the MLE is a consequence of the assumed prior. We note that our focus is different than many simulation studies, where the aim is to evaluate the discrepancies between estimated and true parameter values. Here, we are strictly interested in unintended consequences of prior specifications and its influence on parametric inference.

We further illustrate this issue by fitting occupancy models to empirical point count data collected by McGarigal and McComb (1995). The authors visited over 1,000 sites in the Oregon Coast Range, USA, four times between 1990 and 1992. We fit the basic occupancy model with covariates to detections for three species: gray jay (*Perisoreus Canadensis*), Steller’s jay (*Cyanocitta stelleri*) and song sparrow (*Melospiza melodia*) with two standardized covariates, one representing the distance to forest edges, and the other representing the proportion of the area within 1000 m comprised of mature forest (derived from a gradient nearest neighbor method; Ohmann and Gregory 2002). We specified a uniform prior on detection probability (p ~ Uniform(0,1)), and a normally distributed prior on the intercept and coefficients of occupancy probability with a mean of 0 and *σ*^2^ ranging between 1 and 1000 (1, 10, 100, and 1000). We standardized both covariates, and fit the Bayesian occupancy model with 10,000 MCMC iterations, dropping the first 5,000 as burn-in. Diagnosis of convergence followed the same procedure outlined above for the simulated datasets. We also fit each model in MARK, using the ‘RMark’ package.

To assess the relative prevalence of this issue in the ecological literature we performed a review of the use of priors in Bayesian occupancy modeling. We searched for articles published since 2010 using the term “Bayesian occupancy model” on Web of Science (http://apps.webofknowledge.com). We filtered results to include only those articles published in the field of ecology. We further eliminated any articles that focused on the development and refinement of methods for fitting occupancy. We reviewed a random sample of 25 of the remaining articles, attempting to identify the priors specified

## Results

For the simulated datasets, when the prior standard deviation was small (i.e.σ < 2) the posterior mode was always smaller than the MLE (Fig. 2), as these priors pulled the posterior towards a probability of 0.5. With a standard deviation of 2, the posterior mode was approximately the MLE. At intermediate values of the standard deviation (between 5 and 10), the posterior mode was close to the MLE, but the proximity was influenced by the number of surveyed sites (Fig. 3). As *σ* became large (>100), the posterior became bimodal, with one mode close to the MLE and the other close to 1 (Fig. 2).Importantly, having a large number of sampled sites only buffered against the influence of the prior in a relatively narrow band of values. Generally, the nature ofthe influence of the prior on the posterior and subsequent ecological inference depends on a combination of effects including: 1) the true underlying detection and occupancy probabilities; 2) the number of sampled sites; 3) the number of surveys per site; and, 4) the linear combination of coefficients and covariates. Importantly, the linear combination (i.e. *α* + ***x_i_β***^′^) is the quantity that is transformed, and thus in some cases very large magnitude values for coefficients, when combined with certain values of covariates, could lead to scenarios where the transformation is not impactful (e.g. when there is a strong effect of a covariate that ranges over a very small set of values). However, in other cases, the use of Normal prior distributions with a large *σ* can seriously affect parametric estimates

**Figure 2.**
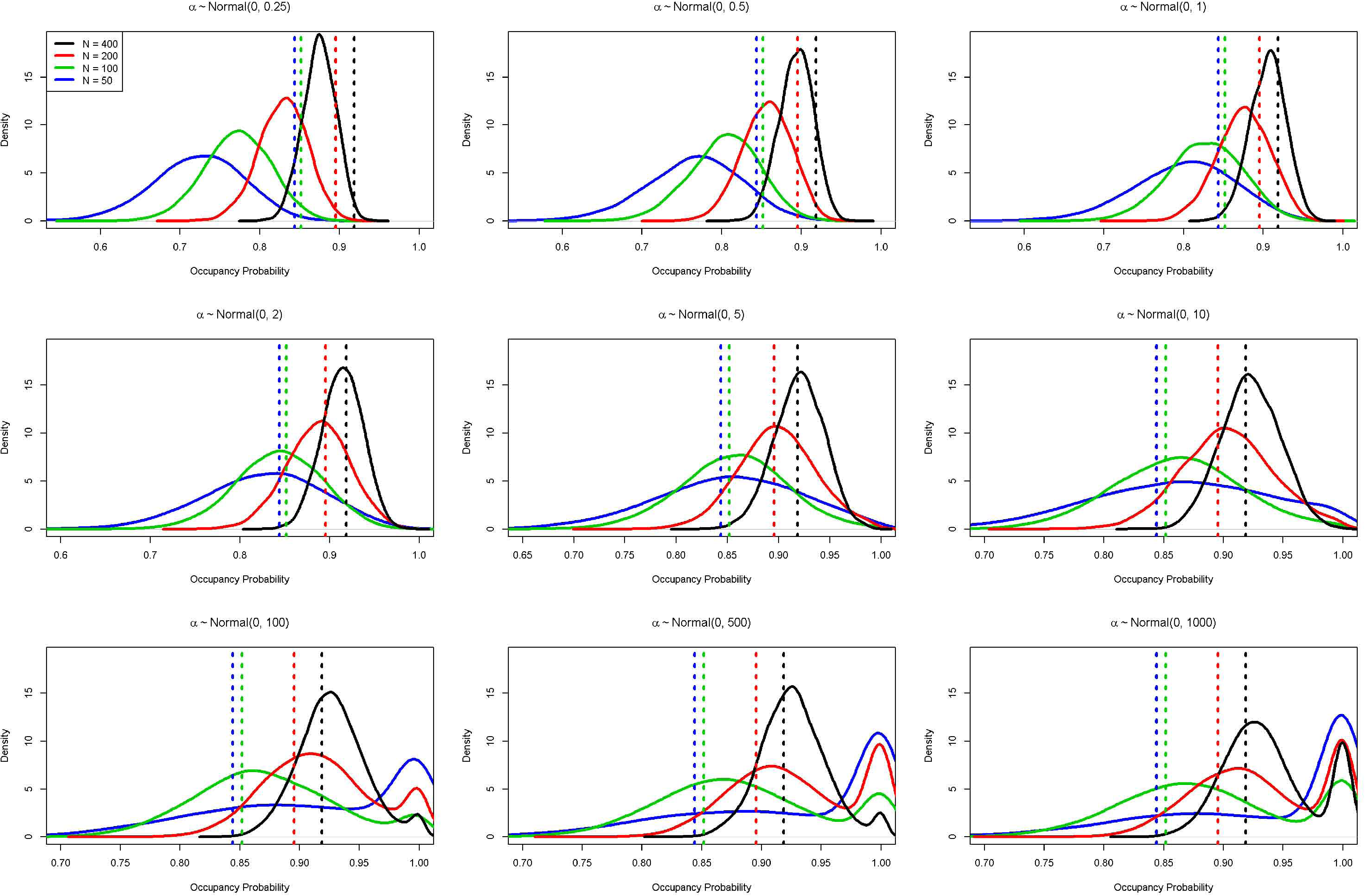
Posterior distributions (solid curves) and the corresponding maximum likelihood estimates (vertical lines) of occupancy probability from simulated data sets of varying number of sites (N = 50, 100, 200, and 400); data were simulated with a true occupancy of 0.9, a per occasion detection probability of 0.2 and 10 sampling occasions. If prior distributions were truly uninformative, the posterior mode would correspond to the maximum likelihood estimate.

In our empirical analysis, there were clear effects of the prior for the gray jay and Steller’s jay data, but not the song sparrow. For both jay species, the median and upper credible bound increased with the prior variance (*σ*^2^), while for the song sparrow, there were no apparent effects of the prior on the posterior (Table S1). For the jays, estimates approaching the MLE could only be obtained by fine-tuning the prior iteratively (results not shown here). Thus, for these datasets, there are very few specifications of the prior distribution that will not impact the posterior distribution and thus our inference on occupancy. Further, many more MCMC iterations were needed to achieve convergence for the gray jay model fit with values of σ^2^ greater than 10, likely due to bimodality similar to that seen in the analysis of simulated datasets (Fig. S3). In addition to causing issues with convergence, the gray jay results also highlight how these priors can impact inference on the habitat factors influencing occupancy. Whether or not credible intervals include 0 is often taken as evidence for the existence of an effect of a covariate on occupancy. For Steller’s jays, the 95% credible intervals (and 95% confidence intervals for the MARK analysis) for the effect of mature forest and edges all overlapped 0, indicating weak evidence for an effect. However, for gray jays the credible intervals (and confidence intervals forthe MARK analysis) did not overlap 0 except in the Bayesian analysis σ^2^ when was set to 1000(Table S1). Thus, we would draw very different conclusions about the influence of mature forest on this species depending only on the specification of the prior. We note here that the low detection probability for gray jays and Steller’s jays could be a result of a lack of closure (i.e. that they were not always available for detection during a survey); however occupancy models are routinely fit to datasets with similar violations of assumptions (see Rota et al. 2009 for adiscussion), and with even lower detection probabilities, and thus we believe this example is still illustrative of the issues that can arise from using a prior that is assumed to be non-informative.

The use of priors that could lead to inferential issues such as those outlined above was common in the recent literature. We found 108 articles published since 2010 that contained the keyphrase “Bayesian occupancy model.” Of the 25 articles reviewed, 8 (approximately one third) reported using priors on α, above, that were incidentally informative on the probability scale. How informative these priors were varied, with some researchers using only moderately informative priors (e.g., uniform distribution between −8 and 8), and others using highly informative priors (e.g., Normal with a variance of >1 000 000). Further, 3 of the articles did not report their priors or described them only as uninformative. Only 2 articles used priors that were likely to be uninformative, based on our results above. We note, that some researchers did report conducting sensitivity analyses to their priors.

### Discussion and suggested guidance

The results that we present above, combined with the apparent prevalence of this issue in the literature, raise concerns about the inference made in regards to species-habitat relationship and resulting distribution patterns. Our literature review, though relatively basic, 249 indicates that this issue might be widespread. Further, most species in a given area are rare (Preston 1948),meaning that researchers likely are fitting models for species with little data (though this depends on the size of the sampling site relative to the species distribution), which will allow for priors to be more influential. However, the true magnitude of the issue is unknown because the circumstances that allow a seemingly uninformative prior distribution to be in fact informative, can vary, depending on the data. As illustrated in our empirical example, there are scenarios under which the specification of the prior will not impact inference; however, there also will be times when specifying a prior that does not impact inference will be difficult and require iterative model fitting. The potential implications of this issue for conservation and management-based studies are significant. We note that camera trapping and the use of occupancy models has become common for studying rare or cryptic, threatened and endangered species (O’Connell et al. 2010). In these studies, both sample sizes and detection probabilities tend to be low, two aspects that can lead to potential issues with influential priors. Even small overestimates in occupancy for such species can have major implications for conservation and management action.

It is important to point out that the highlighted issue is not an inherent shortcoming of Bayesian inference. As a rule, Bayesian analysis requires the specification of priors, and as such, inference will be influenced to some degree by these priors. The models fit above are behaving appropriately and in the above we compared the posteriors to the MLE to illustrate how the priors are influencing results. Priors with large standard deviations or small precisions (inverse of the variance) can strongly influence the posterior distribution of occupancy, both in terms of the occupancy model or just the occupancy parameter, but applies more generally to using a logit Normal prior distribution with a large standard deviation.

The issues outlined above bring up a larger philosophical issue of what type of inference researchers want and why they choose to use Bayesian inference. In many cases, integrating informative prior information with new data to update the belief about an ecological process is not only justifiable but philosophically appealing (Hobbs and Hooten 2015). Informative priors can be particularly useful for the analysis of repeated studies, when there is a desire to include information from published research in current analyses, or to borrow strength across data sources to improve estimate precision (Gopalaswamy et al. 2012). Additionally, informative priors can guard against spurious effects (Gelman et al. 2012, Northrup et al. 2014) and erroneous estimation of large effects in underpowered studies (Lemoine et al. 2016), thus providing more conservative inference than frequentists analyses. But also more generally, informative priors and their shrinkage properties provide a coherent form of model selection (i.e. statistical regularization; Friedman et al. 2001), which has predictive benefits (Gerber et al.2015). However, based on our reading of the literature, many ecologists want inference that is free from effects of the prior (e.g. Jones et al. 2011, Northrup et al. 2015). If researchers truly want inference entirely free from any potential influence of the prior, then they likely should look to different frameworks than Bayesian inference (e.g. using likelihood based occupancy models, or for more complex hierarchical models, methods such as data cloning; Lele et al.2007). If researchers want to obtain Bayesian inference, it is important that they assess the sensitivity of their models to their priors so that they can obtain an understanding of the influence of the prior on their inference. We suggest following a similar approach to the one outlined in our empirical analysis, where sequentially smaller values of σ are used, and posterior medians and credible intervals are compared so that researchers obtain such an understanding. This information can then be used to make an informed decision about the degree to which ecological inference is impacted. For those who desire Bayesian inference and are interested in investigating specific scenarios (i.e.,*ψ*, p, n, number of sites, and *σ*) we provide an easily accessible online tool for doing so (https://briangerber.shinyapps.io/OccupancyPrior/).

As noted above, simply using a more restrictive prior might not provide the desired inference in all scenarios. The literature offers some further guidance on priors for logistic regression and occupancy. Gelman et al. (2008) suggest the use of a Cauchy distribution with center 0 and scale 2.5 as a default prior when conducting logistic regression. However, this prior still displays slight bimodality at 0 and 1 and thus has the potential to affect posterior distributions. An exact prior distribution that is completely invariant to transformation, such as between the logit and probability scale, is the Jeffry’s prior (Jeffreys 1946), which can be derived for occupancy models, but also will likely be informative and thus might not be more appropriate than a Normal distribution with small variance. We note that recent publications in statistical ecology describing how to fit occupancy models in a Bayesian framework provide some suggestions for priors that should reduce the concerns we raise here. Kéry and Royle (2015), in worked examples suggest, for the intercept, a uniform distribution between 0 and 1 that is then logit-transformed, though with unscaled covariates this prior could lead to estimates of the intercept that are biased low. Further, the uniform distribution actually leads to an improper posterior, though the degree to which this impacts inference is likely limited. Following our findings and the recommendation from Hobbs and Hooten (2015), logit-Normal prior distributions with a variance near 2 (standard deviation approximately 1.4) will often likely be weakly informative (when covariates are standardized), but we still strongly suggest conducting a prior sensitivity analysis. We also note, that while this issue is relatively well understood in the types statistical ecology texts discussed above, these texts often are unapproachable for practitioners hoping to apply models.

Beyond the above suggestions, practitioners should use particular care when detection is low, few sites are surveyed, and occupancy is very low or very high. Generally, caution should be applied when parameter uncertainty is likely to be large and near the probability boundaries, zero and one. In all cases, but particularly under these circumstances, we suggest first scaling all independent variables so that large magnitude coefficient estimates are avoided (Gelman and Hill 2007; scaling also speeds convergence in many cases). Secondly, we suggest fitting models with a range of priors, to assess their influence. We caution that surveying many sites is not a panacea for this issue. In the dataset analyzed above, there were 4 visits to over 1,000 sites, a dataset that dwarfs most used in occupancy analyses.

## Conclusion

Complex computational and statistical methods will continue to become more attainable for ecologists and other practitioners as computers become more advanced, and books are published that provide walk-through examples and code to fit complicated models. While the issue of uninformative priors becoming informative when transformed is well known to statisticians (Lele and Dennis 2009, Seaman III et al. 2012), many of the previous descriptions of this problem are unapproachable for ecologists. We hope that this comment will spur other ecologists to take care to better understand the models that they fit, and what their model outputs and results and the associated ecological inference truly means. The tools are available for us to fit difficult and complex models, the onus is thus on us to understand what they mean.

## Acknowledgements

The authors thank K. McGarigal for use of the empirical data presented. M.G. Betts, B.R. Noon, L. BaileyE, B.A. MosherE, and D. Martin provided comments on earlier drafts that greatly improved the manuscript.

## Supplementary material

Figure S1: Depiction of transformation between logit and probability scale

Figure S2: Animation of relationship between normal prior distributions on logit and probability scales

Figure S3: Supplemental results for Gray Jay analysis

Table S1: Supplemental results for analysis of avian occupancy

Appendix S1: Example JAGS code to fit occupancy model with covariates

Appendix S2: Example JAGS code to fit occupancy model with no covariates

## References

Clark, J.S, (2005) Why environmental scientists are becoming Bayesians. Ecology Letter 8: 2–14.

Elith, J., J.R., Leathwick (2009) Species distribution models: ecological explanation and prediction across space and time. Annual Review of Ecology, Evolution, and Systematics 40: 677.

Fiske, I., and R. Chandler. (2011) unmarked: An R package for fitting hierarchical models of wildlife occurrence and abundance. Journal of Statistical Software 43: 1–23.

Friedman, J., T. Hastie, and R. Tibshirani. 2001. The elements of statistical learning. Springer series in statistics Springer, Berlin.

Gelman, A., J.B. Carlin, H.S. Stern, and D.B. Rubin. 2014. Bayesian data analysis. Chapman & Hall/CRC Boca Raton FL, USA.

Gelman, A., J. Hill. 2007. Data analysis using regression and multilevel/hierarchical models. Cambridge University Press, Cambridge; New York.

Gelman, A., J. Hill, and M. Yajima. 2012. Why We (Usually) Don’t Have to Worry About Multiple Comparisons. Journal of Research on Educational Effectiveness5: 189–211.

Gelman, A., A. Jakulin, and M.G. Pittau, and Y.-S. Su. 2008. A weakly informative default prior distribution for logistic and other regression models. The annals of applied statistics:1360–1383.

Gelman, A. and D.B. Rubin. 1992. Inference from iterative simulation using multiple sequences. Statistical Science 7: 457–511.

Gerber, B.D., W.L. Kendall, M.B. Hooten, J.A. Dubovsky, and R.C. Drewein. 2015. Optimal population prediction of sandhill crane recruitment based on climate-mediated habitat limitations. Journal of Animal Ecology.84: 1299–1310.

Gopalaswamy, A.M., J.A. Royle, M. Delampady, J.D. Nichols, K.U. Karanth, and D.W. Macdonald, 2012. Density estimation in tiger populations: combining information for strong inference Ecology 93:1741–1751

Guisan, A., W. Thuller. 2005. Predicting species distribution: offering more than simple habitat models. Ecology Letters 8: 993–1009

Hines, J. 2006. PRESENCE. Software to estimate patch occupancy and related parameters. USGS, Patuxent Wildlife Research Center, Laurel, Maryland, USA

Hobbs, N.T., and M.B. Hooten. 2015. Bayesian models:a statistical primer for ecologists. Princeton University Press

Jeffreys, H., 1946. An invariant form for the prior probability in estimation problems. Pages 453–461 in Proceedings of the Royal Society of London A: Mathematical, Physical and Engineering Sciences. The Royal Society.

Jones, J.E., A.J. Kroll, J. Giovanini, S.D. Duke, M.G. Betts. 2011. Estimating thresholds in occupancy when species detection is imperfect. Ecology. 92: 2299–2309

Kéry, M., and J.A. Royle. 2015 Applied Hierarchical Modeling in Ecology: Analysis of distribution, abundance and species richness in R and BUGS: Volume 1: Prelude and Static Models Academic Press.

Laake, J.L. 2013. RMark: an R interface for analysis of capture-recapture data with MARK. US Department of Commerce, National Oceanic and Atmospheric Administration, National Marine Fisheries Service, Alaska Fisheries Science Center.

Lele, S.R. and B. Dennis. 2009 Bayesian methods for hierarchical models: Are ecologists making a Faustian bargain. Ecological Applications 19: 581–584.

Lele, S.R., B. Dennis, and F. Lutscher. 2007. Data cloning: easy maximum likelihood estimation for complex ecological models using Bayesian Markov chain Monte Carlo methods. Ecological Letters 10: 551–563.

Lemoine, N.P., A. Hoffman, A.J. Felton, L. Baur, F. Chaves, J. Gray, Q. Yu, and M.D. Smith. 2016. Underappreciated problems of low replication in ecological field studies. Ecology 97: 2554–2561.

MacKenzie, D.I., J.D. Nichols, G.B. Lachman, S. Dreoge, J. Andrew Royale, and C.A. Langtimm, 2002 Estimating site occupancy rates when detection probabilities are less than one. Ecology 83: 2284–2255

McCarthy, M.A. 2007. Bayesian methods for ecology. Cambridge University Press

McGarigal, K., and W.C. McComb. 1995. Relationships between landscape structure and breeding birds in the Oregon Coast Range. Ecological monographs 65:235–260.

Northrup, J.M., C.R. Anderson Jr., and G. Wittemyer. 2015. Quantifying spatial habitat loss from hydrocarbon development through assessing habitat selection patterns of mule deer Global change biology 21: 3961–3970

Northrup, J.M., A.B.A. Shafer, C.R. Anderson, D.W. Coltman, and G. Wittemyer. 2014. Fine-scale genetic correlates to condition and migration in a wild cervid Evolutionary Applications 7: 937–948.

O’Connell, A. F., J. D. Nichols, and K. U. Karanth. 2010. Camera traps in animal ecology:methods and analyses. Springer Science & Business Media.

Ohmann, J. L., and M. J. Gregory. 2002. Predictive mapping of forest composition and structure with direct gradient analysis and nearest-neighbor imputation in coastal Oregon, U.S.A. Canadian Journal of Forest Research 32: 725–741

Plummer, M. 2015a. JAGS Version 4.0.0. user manual.

Plummer, M. 2015b. rjags: Bayesian graphical models using MCMC.R. package version 3-15. http://CRAN.R-project.org/package=rjags.

Preston, F.W. 1948 The Commonness, And Rarity, of Species. Ecology.29: 254–283.

Rota, C.T., R.J. Fletcher Jr., R.M. Dorazio, and M.G. Betts. 2009. Occupancy estimation and the closure assumption. Journal of Applied Ecology 46: 1173–1181.

Royle, J.A., and R.M. Dorazio. 2008. Hierarchical modeling and inference in ecology: the analysis of data from populations, metapopulations and communities. Academic Press.

Seaman, J.W. III, J.W. Seaman Jr, and J.D. Stamey. 2012. Hidden dangers of specifying noninformative priors. The American Statistician 66: 77–84.

Team, R.C. 2015. R: A language and environment for statistical computing. R Foundation for Statistical Computing, Vienna, Austria. http://www.R-project.org/.

White, G.C., and K.P. Burnham. 1999. Program MARK: survival estimation from populations of marked animals. Bird Study 46: S120–S139.

